# Sensitive detection of pre-integration intermediates of LTR retrotransposons in crop plants

**DOI:** 10.1101/317479

**Authors:** Jungnam Cho, Matthias Benoit, Marco Catoni, Hajk-Georg Drost, Anna Brestovitsky, Matthijs Oosterbeek, Jerzy Paszkowski

**Affiliations:** The Sainsbury Laboratory, University of Cambridge, Cambridge CB2 1LR, UK

## Abstract

Retrotransposons have played an important role in the evolution of host genomes^1,2^. Their impact on host chromosomes is mainly deduced from the composition of DNA sequences, which have been fixed over evolutionary time. These studies provide important “snapshots” reflecting historical activities of transposons but do not predict current transposition potential. We previously reported Sequence-Independent Retrotransposon Trapping (SIRT) as a methodology that, by identification of extrachromosomal linear DNA (eclDNA), revealed the presence of active LTR retrotransposons in *Arabidopsis*^9^. Unfortunately, SIRT cannot be applied to large and transposon-rich genomes of crop plants. We have since developed an alternative approach named ALE-seq (*<u>a</u>*mplification of *<u>L</u>*TR of *<u>e</u>*clDNAs followed by *<u>seq</u>*uencing). ALE-seq reveals sequences of 5’ LTRs of eclDNAs after two-step amplification: *in vitro* transcription and subsequent reverse transcription. Using ALE-seq in rice, we detected eclDNAs for a novel *Copia* family LTR retrotransposon, *Go-on*, which is activated by heat stress. Sequencing of rice accessions revealed that *Go-on* has preferentially accumulated in *indica* rice grown at higher temperatures. Furthermore, ALE-seq applied to tomato fruits identified a developmentally regulated *Gypsy* family of retrotransposons. Importantly, a bioinformatic pipeline adapted for ALE-seq data analyses allows the direct and reference-free annotation of new active retroelements. This pipeline allows assessment of LTR retrotransposon activities in organisms for which genomic sequences and/or reference genomes are unavailable or are of low quality.

---

Chromosomal copies of activated retrotransposons containing long terminal repeats (LTRs) are transcribed by RNA polymerase II, followed by reverse transcription of transcripts to extrachromosomal linear DNAs (eclDNA); these integrate back into host chromosomes. Because of the two obligatory template switches during reverse transcription, the newly synthetized eclDNA is flanked by LTRs of identical sequence. Their subsequent divergence due to the accumulation of mutations correlates well with length of time since the last transposition, and thus transposon age^3^. However, the age of LTR retrotransposons cannot be used to predict their current transpositional potential. Moreover, predictions are further complicated by recombination events that occur with high frequency between young and old members of a retrotransposon family^4^; thus old family members also contribute to the formation of novel recombinant elements that insert into new chromosomal positions. Although, retrotransposon activities can be easily measured at the transcriptional level^5,6^, the presence of transcripts is a poor predictor of transpositional potential due to posttranscriptional control of this process^7,8^. In addition, direct detection of transposition by genome-wide sequencing to identify new insertions is too expensive and time-consuming to be applied as a screening method. Clearly, the development of an expeditious approach to identify active retrotransposons that predict their transposition potential would be welcomed. We previously described the SIRT strategy for *Arabidopsis* that led to the identification of eclDNA of a novel retroelement and subsequent detection of new insertions^9^. Thus, the presence of eclDNAs, the last pre-integration intermediate, was shown to be a good predictor of retrotransposition potential.

Retrotransposons include a conserved sequence known as the primer binding site (PBS), where binding of the 3’ end of cognate tRNA initiates the reverse transcription reaction^9^. Met-iCAT PBS was chosen for SIRT as it is the site involved in the majority of annotated *Arabidopsis* retrotransposons^9^. To examine whether Met-iCAT PBS sequences are also predominant in LTR retrotransposons of other plants, we used the custom-made software *LTRpred* for *de novo* annotation of LTR retrotransposons in rice and tomato genomes (see Materials and methods). Young retroelements were selected by filtering for at least 95% identity between the two LTRs and subsequently examined for their cognate tRNAs (Supplementary Figure 1a). As in *Arabidopsis*, around 80% of LTR retrotransposons in the tomato genome contained Met-iCAT PBS (Supplementary Figure 1a). In contrast, only 30% harboured Met-iCAT PBS in rice, and Arg-CCT PBS was involved in 60% of young LTR retrotransposons (Supplementary Figure 1a). Nonetheless, we used Met-iCAT PBS in our initial experiments because most retrotransposons known to be active in rice callus (e.g. *Tos17* and *Tos19*) contain Met-iCAT PBS. Initially, SIRT was performed on DNA extracted from rice leaves and callus; however, we did not detect eclDNAs for *Tos17* and *Tos19* in rice tissues by this method (Supplementary Figure 2a and b). We reasoned that the short stretch of PBS used for primer design in SIRT may have impaired PCR efficiency due to the many PBS-related sequences present in larger genomes containing a high number of retroelements, as is the case in rice.

To counter this problem, we developed an alternative method, named *<u>a</u>*mplification of *<u>L</u>*TR of *<u>e</u>*clDNAs followed by *<u>seq</u>*uencing (ALE-seq), with significantly improved selectivity and sensitivity of eclDNA detection. A crucial difference to SIRT is that ALE-seq amplification of eclDNA is separated into two reactions: *in vitro* transcription and reverse transcription (Figure 1a). This decoupling of the use of the two priming sequences by production of an RNA intermediate is significantly more selective and efficient than the single PCR amplification in SIRT.

**Figure 1.**
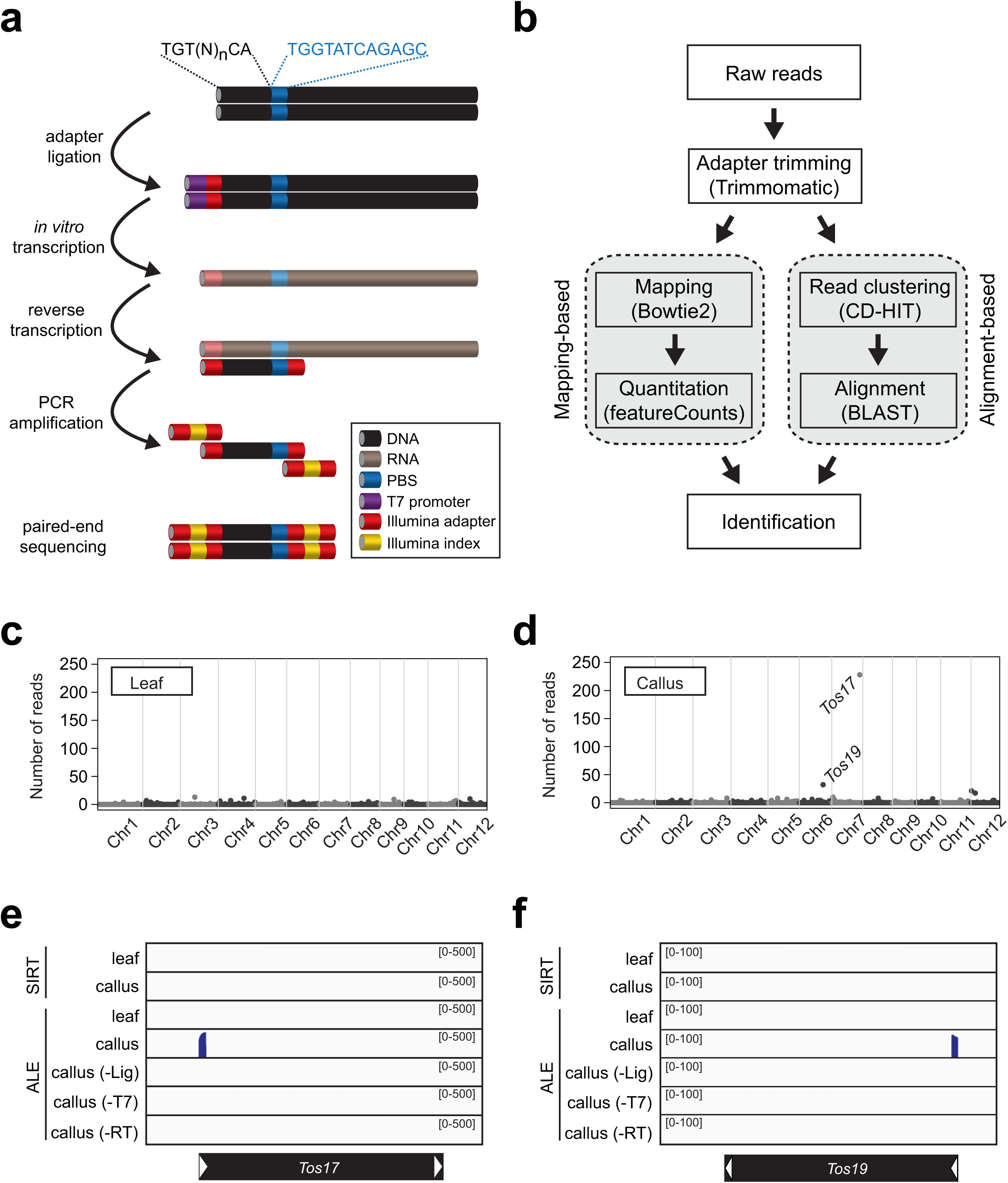
Detection of eclDNA by ALE-seq. **a**, The workflow of ALE-seq. The colour code is indicated in a box. **b**, Analysis pipeline of ALE-seq results. The sequenced reads can be mapped to the reference genome or aligned to each other to obtain a cluster consensus. **c** and **d**, Genome-wide plots of rice ALE-seq results from leaf (**c**) and callus (**d**). The levels are shown as number of reads mapped to each retrotransposon. Dots represent annotated retrotransposons; those corresponding to *Tos17* and *Tos19* are indicated. **e** and **f**, Read plots mapped to *Tos17* (**e**) and *Tos19* (**f**). The black bars represent retrotransposons and white arrowheads indicate LTRs.

ALE-seq starts with ligation to the ends of eclDNA of an adapter containing a T7 promoter sequence at its 5’ end and subsequent *in vitro* transcription with T7 RNA polymerase. The synthesized RNA is then reverse transcribed using the primer that binds the transcripts at the PBS site. The adapter and the oligonucleotides priming reverse transcription are anchored with partial Illumina adapter sequences, which allows the amplified products to be directly deep-sequenced in a strand-specific manner. The ALE-seq-sequences derived retrotransposon eclDNAs are predicted to contain the intact 5’ LTR up to the PBS site, flanked by Illumina paired-end sequencing adapters. We used the Illumina Mi-seq platform for sequencing because its long reads of 300 bp from both ends cover the entire LTR lengths of most potentially active elements. It is worth noting that the Illumina adapters were tagged to the intact LTR DNA without fragmentation of the amplicons. This together with the long reads of Mi-seq allowed us to reconstitute the complete LTR sequences, even in the absence of the reference genome sequence. The reconstituted LTRs were analysed using the alignment-based approach that complements the mapping-based approach when the reference genome is incomplete (Figure 1b).

First, we tested ALE-seq on *Arabidopsis* by examining heat-stressed Col-0 *Arabidopsis* plants^11^, *met1-1* mutant^9^ and epi12^12^, a *met1-*derived epigenetic recombinant inbred line. ALE-seq cleanly and precisely recovered sequences of complete LTRs for *Onsen*, *Copia21* and *Evade* in samples containing their respective eclDNA (Supplementary Figure 3a to g)^8,9,11^. Due to priming of the reverse transcription reaction at PBS, the reads were explicitly mapped to the 5’ but not to the 3’ LTR, although the two LTRs have identical sequences. The ALE-seq reads have well-defined extremities, starting at the position marking the start of LTRs and finishing at the PBS, which is consistent with their eclDNA origin. The ends of LTRs can also be inspected for conserved sequences that would further confirm their eclDNA origin (Supplementary Figure 1b). This reduced ambiguity of read mapping in ALE-seq analysis, combined with the clear-cut detection of LTR ends, allows for explicit and precise assignment of ALE-seq results to active LTR retrotransposons.

Since SIRT failed to detect eclDNAs of rice retrotransposons known to be activated in rice callus, we examined whether ALE-seq would identify their eclDNAs. As shown in Figure 1c to f, ALE-seq unambiguously detected eclDNAs of *Tos17* and *Tos19* in rice callus, but not in leaf samples. To test whether detection of 5’ LTR sequences requires the entire ALE-seq procedure, we performed control experiments with depleted ALE-seq reactions, for example, in the absence of enzymes for either ligation, *in vitro* transcription, or reverse transcription. All incomplete procedures failed to produce sequences containing 5’ LTRs derived from eclDNAs (Figure 1e and f). Taken together, the data show that ALE-seq can detect eclDNAs of LTR retrotransposons in *Arabidopsis* as well as in rice with considerably greater efficiency than the SIRT method.

To examine the suitability of ALE-seq for quantitative determination of eclDNA levels, we carried out a reconstruction experiment spiking 100 ng of genomic DNA from rice callus with differing amounts of PCR-amplified full-length *Onsen* DNA from 1 ng to 100 fg (Figure 2a to d). The results in Figure 2a and b show that the readouts of ALE-seq for *Onsen* correlate well with the input amounts (R^2^=0.99). The initial ALE-seq steps of ligation and *in vitro* transcription impinged proportionally on the input DNA, resulting in unbiased quantification of the eclDNA and minimal quantitative distortion of the final ALE-seq data. Noticeably, the levels of *Tos17* were similar in all the spiked samples, indicating that addition of *Onsen* DNA did not influence the detection sensitivity of *Tos17*, at least for the amounts tested (Figure 2c and d). Thus, ALE-seq can be used to accurately determine eclDNA levels.

**Figure 2.**
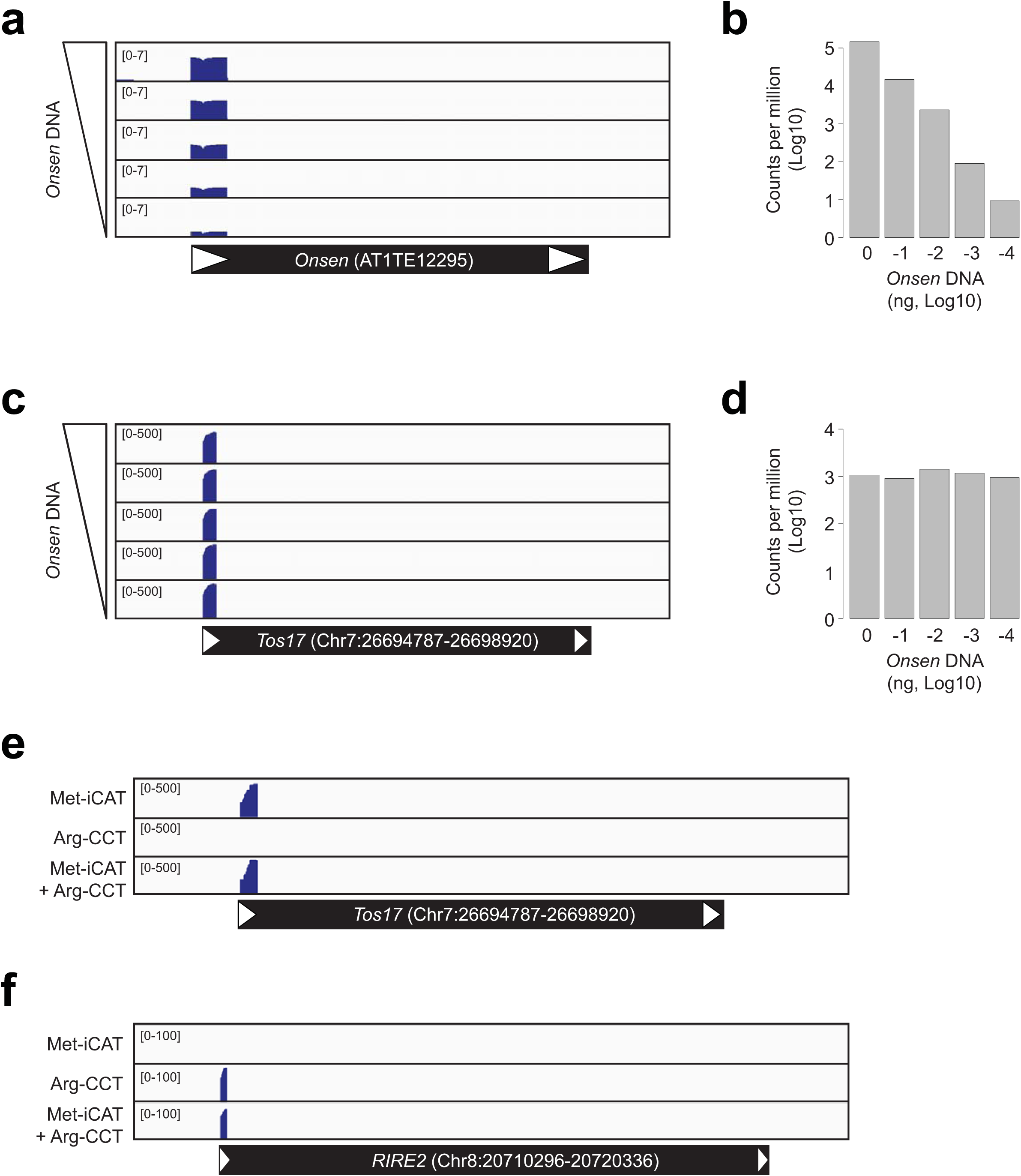
Sensitivity and specificity of eclDNA detection by ALE-seq. **a**-**d**, ALE-seq reconstruction experiment with varying amounts of PCR-amplified *Onsen* DNA added to rice callus DNA. Genome browser image with the read coverage (**a** and **c**) and quantitated read counts (**b** and **d**) for *Onsen* (**a** and **b**) and *Tos17* (**c** and **d**) loci. The amounts of *Onsen* DNA added were 1 ng, 100 pg, 10 pg, 1 pg or 100 fg; 100 ng of rice callus DNA was used. Note that read coverage values are Log10-converted in **a**. For **b** and **d**, values are shown as Log10-converted counts per million reads. **e** and **f**, Read coverage plots for the ALE-seq of rice callus using different RT primers. *Tos17* and *RIRE2* transposons depicted below the plots as in Figure 1.

Most rice retrotransposons harbour Arg-CCT PBS (Supplementary Figure 1a). We tested whether the reverse transcription reaction can be multiplexed to capture both types of retrotransposons (containing Arg-CCT or Met-iCAT PBS) and whether multiplexing of the reverse transcription primers compromises the sensitivity of the procedure. ALE-seq was performed on DNA from rice callus, testing each of the reverse transcription primers separately or as a mixture of both primers in a single reaction. As shown in Figure 2e, the levels of *Tos17* recorded in the samples with both primers were similar to the Met-iCAT primer alone. Importantly, we also detected the eclDNAs of the *RIRE2* element containing Arg-CCT PBS (Figure 2f), which was known to be transpositionally active in rice callus^7^.

We next used ALE-seq to search for novel active rice retrotransposons. Since many plant retrotransposons are transcriptionally activated by abiotic stresses^11,13^, we subjected rice plants to heat stress before subjecting them to ALE-seq. In this way we identified a *Copia*-type retrotransposon able to synthetize eclDNA in the heat-stressed plants (Figure 3a to c) and named this element *Go-on* (the Korean for ‘high temperature’). The three retrotransposons with the highest eclDNA levels in heat-stress conditions all belong to the *Go-on* family (Figure 3b and Supplementary Figure 4a). Although, eclDNAs were detected for all three copies, *Go-on3* seems to be the youngest and, thus, possibly the most active family member, containing identical LTRs and a complete ORF (Supplementary Figure 4a). As depicted in Supplementary Figure 4a, the 5’ LTR sequences of the three *Go-on* copies are identical; thus the ALE-seq reads derived from *Go-on3* LTR were also cross-mapped to other copies that are possibly inactive or have reduced activities. To further determine whether sequences of *Go-on* LTRs recovered by ALE-seq are indeed derived from *Go-on3* or also from other family members, we performed an ALE-seq experiment using RT primers located further downstream of the PBS, including sequences specific for each *Go-on* family member (Supplementary Figure 4a). The amplified ALE-seq products revealed that the eclDNAs produced in heat-stressed rice originated only from *Go-on3* (Supplementary Figure 4b). We validated the production of eclDNAs of *Go-on3* by sequencing the junction of the adapter and the 5’ end of LTR (Supplementary Figure 4c) and by qPCR (Supplementary Figure 5a).

**Figure 3.**
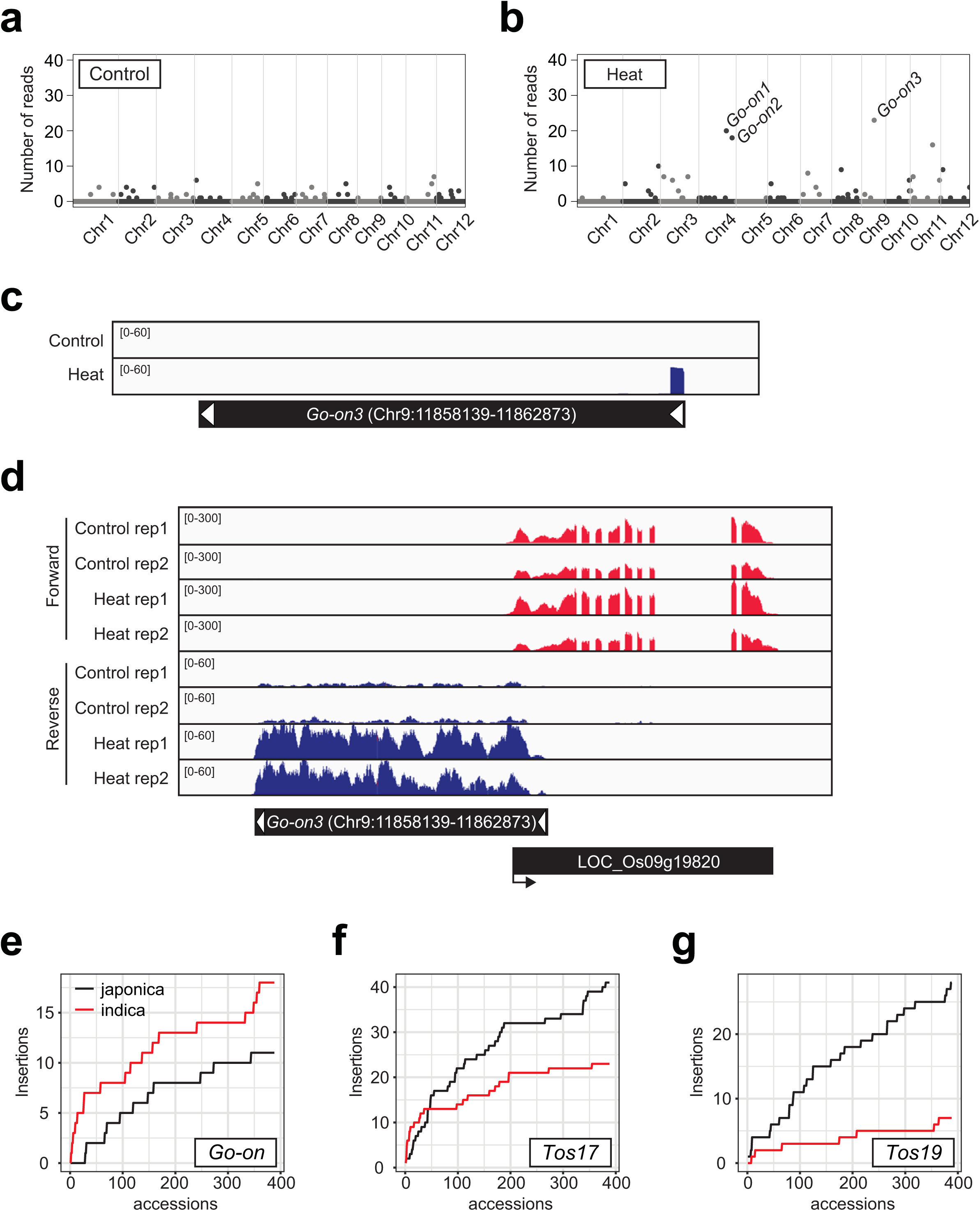
Identification of a novel heat-activated retrotransposon in rice. **a** and **b**, Genome-wide plots of rice ALE-seq results as in Figure 1. Control (**a**) and heat-stressed (**b**) rice plants were used. One-week-old seedlings were subjected to heat stress (44°C) for 3 days. The levels are shown as the number of reads mapped to the retroelements. Three *Go-on* copies are indicated in **b**. **c**, Read coverage plot for *Go-on3*. **d**, RNA-seq data showing *Go-on3* and a neighbouring gene. RNA-seq data were generated using the same plant materials as in **a** and **b**. **e**-**g**, Cumulative plots for the number of unique insertions of *Go-on* (**e**), *Tos17* (**f**), and *Tos19* (**g**) in the genomes of 388 *japonica* and *indica* rice accessions.

Next, we examined whether *Go-on3* is transcriptionally activated in rice subjected to heat stress. RNA-seq and the RT-qPCR data clearly showed that *Go-on* is strongly activated in heat-stress conditions (Figure 3d and Supplementary Figure 5b). The LTR sequence of *Go-on3* contains the heat-responsive sequence motifs (Supplementary Figure 5c), which is consistent with its heat stress-mediated transcriptional activation (Figure 3d). To determine whether *Go-on* is also activated in *indica* rice, we heat-stressed plants of *IR64* for three days and examined *Go-on* RNA and DNA levels. Similar to *japonica* rice, *Go-on* RNA and DNA accumulated markedly under heat stress (Supplementary Figure 6a and b), suggesting that the trigger for *Go-on* activation is conserved in both of these evolutionarily distant rice genotypes. Analysis of the RNA-seq data from the heat-stressed rice plants revealed a poor correlation between the mRNA and eclDNA levels of retrotransposons (Supplementary Figure 7a and b). This agrees with the notion that the eclDNA level is a better predictor of retrotransposition than the RNA level.

To determine whether *Go-on* proliferation increases in rice grown at elevated temperatures, we analysed the historical retrotransposition of *Go-on* using the genome resequencing data of rice accessions from the 3,000 Rice Genome Project^14^. First, we retrieved the raw sequencing data for all 388 *japonica* rice accessions and the same number of randomly selected sequences of *indica* rice accessions. Using the Transposon Insertion Finder (TIF) tool^15^, *japonica* and *indica* sequences were analysed for the number of *Go-on* copies and their genome-wide distribution. Only unique insertions that were absent in the reference genome were scored and the cumulative number of new insertions was plotted (Figure 3e to g). Figure 3e shows that the *indica* rice population grown in a warmer climate accumulated significantly more *Go-on* copies than the *japonica* population. As controls, we also examined the accumulation of *Tos17* and *Tos19*, which were not known to be activated by heat stress. Both retrotransposons showed more transposition events in *japonica* than in *indica* rice (Figure 3f and g). Therefore, *Go-on* as a heat-activated retroelement has undergone specific accumulation in *indica* rice subjected to a warmer climate.

It was reported previously that the tomato genome experiences a significant loss of DNA methylation in fruits during their maturation, which leads to transcriptional activation of retrotransposons^16,17^. However, it was not known whether these transcriptionally activated tomato transposons synthesise eclDNA. It was questionable whether the ALE-seq strategy is sensitive enough to detect eclDNA in the tomato genome, which is three times larger than that of rice^18^. To address these questions, ALE-seq was carried out on DNA samples from fruits at 52 days post anthesis (DPA), when the loss of DNA methylation is most pronounced^16^, and from leaves as a control. It is important to note that we used tomato cultivar (cv.) M82 for these experiments, as it is commonly used for genetic studies^19,20^, and that the sequence of the current tomato reference genome is based on cv. Heinz 1706^18^. Since retrotransposon sequences and their chromosomal distributions differ largely between genomes of different varieties within the same plant species^21–24^, we could not use the standard mapping-based annotation of the ALE-seq results. As a consequence, we developed a reference-free and alignment-based approach that adopts the clustering of reads based on their sequence similarities (Figure 1b). Briefly, the reads from both samples were pooled and then clustered by sequence homology (See Materials and methods). The consensus of each cluster was determined and used as the reference in paired-end mapping. Subsequently, the consensus sequences were used for a BLAST search against the reference genome for the closest homologues. In this way, the BLAST search was able to map the clustered ALE-seq output to reference genome annotated retrotransposons, which are most similar to the ALE-seq recovered sequences. Applying this strategy, we identified a retroelement belonging to a *Gypsy* family (*FIRE, Fruit-Induced RetroElement*) that produces significant amounts of eclDNA at 52 DPA during fruit ripening (Figure 4a). We also determined the transcript levels of the *FIRE* element in leaves and 52 DPA fruit samples. As shown in Figure 4b, fruit RNA levels were enhanced twofold compared to leaves, where *FIRE* eclDNA was barely detectable (Figure 4a). Finally, we found that the DNA methylation status of the *FIRE* element was lower in fruits than leaves in all three sequence contexts (Figure 4c and e). In contrast, the DNA methylation levels of sequences directly flanking *FIRE* were similar in leaves and fruits (Figure 4d to f).

**Figure 4.**
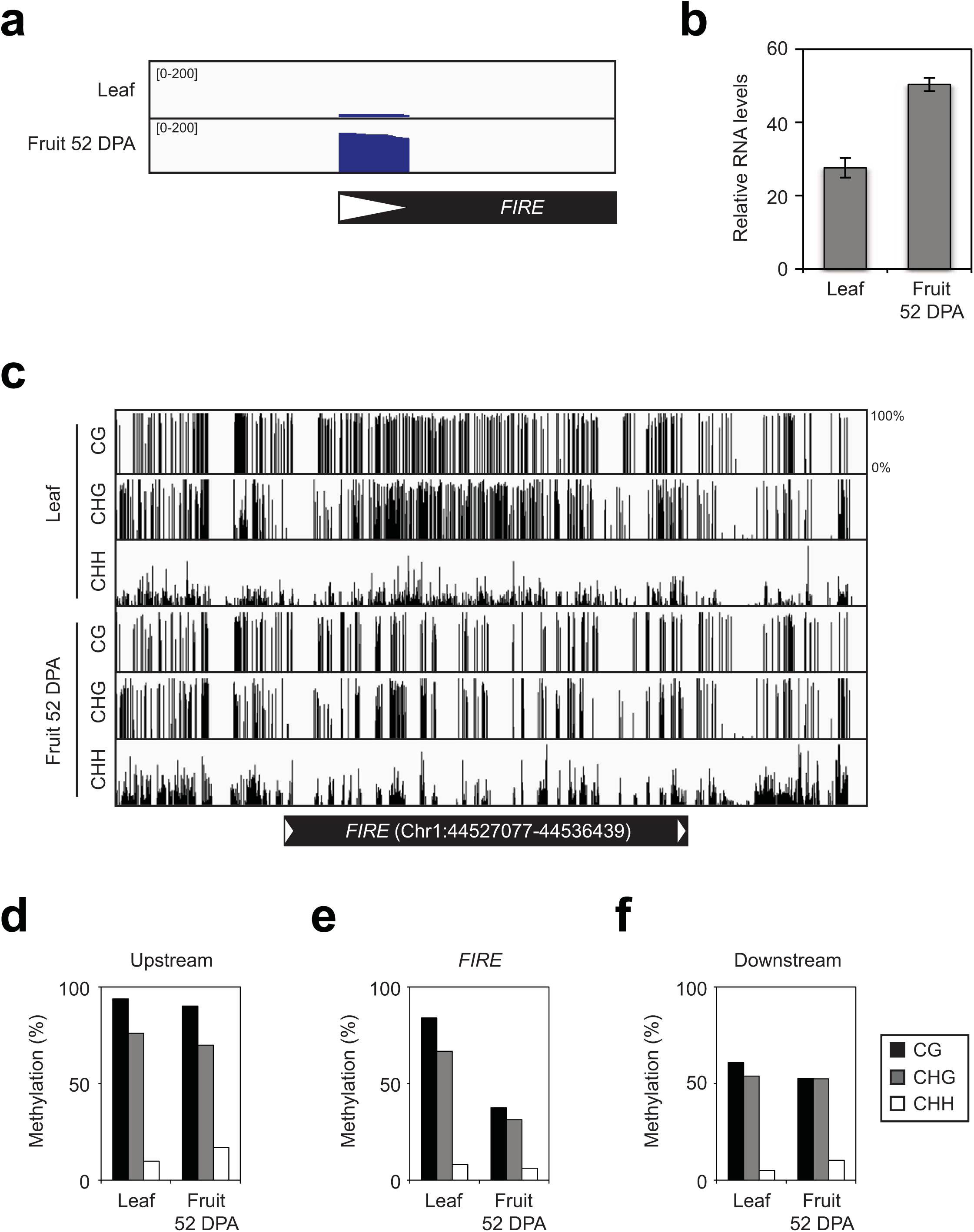
Identification of a tomato retrotransposon activated in fruit pericarp. **a**, Read coverage plot for the *FIRE* retrotransposon identified in tomato fruit pericarp. **b**, The RNA levels of *FIRE* in leaves and fruits determined by RT-qPCR. The levels are means ± standard deviation (sd) of two biological replicates performed with three technical repeats each. Normalization was done against *SlCAC* (Solyc08g006960). **c**, Genome browser image for the DNA methylation levels at *FIRE* elements in leaves and fruits of tomato. **d**-**f**, Quantitation of DNA methylation levels. The levels are the averages of percent DNA methylation in the indicated regions. The upstream and downstream regions are immediate flanking sequences with the same length as *FIRE*.

Recently, a novel active retrotransposon was identified in rice by sequencing extrachromosomal circular DNA (eccDNA) produced as a by-product of retrotransposition or by nuclear recombination reactions of eclDNAs^25–27^. Although the method of eccDNA sequencing has certain advantages over SIRT, such as increased sensitivity and the recovery of sequences of the entire element, it also has certain limitations. For example, the method requires relatively large amounts of starting material but still shows serious limits in sensitivity and indicative power for retrotransposition. The method did not detect the eccDNA of *Tos19* in rice callus, where this transposon is known to move^25^. Most importantly, eccDNAs may also be the result of genomic DNA recombination^27^ and these background products may be misleading when extrapolating to the transpositional potential of a previously unknown element. In this respect, ALE-seq is a significantly improved tool that largely overcomes the above-mentioned limitations of previous methods, and requires only 100 ng of plant DNA.

The heat-responsiveness of *Go-on*, the novel heat-activated *Copia* family retrotransposon of rice detected using ALE-seq, seems to be conferred by *cis*-acting DNA elements embedded in the LTR, which are similar to the heat-activated *Onsen* retrotransposon in *Arabidopsis*^11^. Although heat stress can induce production of mRNA and eclDNA of *Onsen*, its retrotransposition is tightly controlled by the small interfering RNA pathway^11^. Given that real-time transposition of rice retrotransposons has only been detected in epigenetic mutants^28,29^ and triggered by tissue culture conditions causing vast alterations in the epigenome^7,30^,or as a result of interspecific hybridization^31^, an altered epigenomic status seems to be an important prerequisite for retrotransposition. In fact, we failed to detect transposed copies of *Go-on* in the progeny of heat-stressed rice plants. Thus, although *Go-on* produces eclDNAs after heat stress, it may be mobilized only at low frequency in wild type rice due to epigenetic restriction of retrotransposition. Nevertheless, on an evolutionary scale, the higher copy number of *Go-on* in *indica* rice populations grown at elevated temperatures is compatible with potential mobility.

Many retrotransposons are transcriptionally reactivated during specific developmental stages or in particular cell types^32–34^. In tomato, fruit pericarp exhibits a reduction in DNA methylation during ripening. This is largely attributed to higher transcription of the *DEMETER-LIKE2* DNA glycosylase gene^17,35–37^. Despite massive transcriptional reactivation of retrotransposons in tomato fruits, it has been difficult to determine whether further steps toward transposition also take place. Using ALE-seq, we identified eclDNA that we annotated using a reference-free and alignment-based approach to a novel *FIRE* element. *FIRE* has 164 copies in the reference tomato genome and in a conventional mapping-based approach the ALE-seq reads of *FIRE* cross-mapped to multiple copies, making it difficult to assign eclDNA levels to particular family members. Therefore, our annotation strategy can be used in situations where sequence of the reference genome is unavailable or the mapping of reads is hindered by the high complexity and multiplicity of the retrotransposon population.

ALE-seq could also be applied to non-plant systems. For example, numerous studies in various eukaryotes, including mammals, found that retrotransposons are transcriptionally activated by certain diseases or at particular stages during embryo development^38–40^. It was also suggested that retrotransposition might be an important component of disease progression^41^. Given that the direct detection of retrotransposition is challenging, it would be interesting to use ALE-seq to determine whether such temporal relaxations of epigenetic transposon silencing also result in the production of the eclDNAs, as the direct precursor of the chromosomal integration of a retrotransposon.

## Materials and methods

### Plant materials

Seeds of *Oryza sativa ssp. japonica cv. Nipponbare* and *Oryza sativa ssp. indica cv. IR64* were surface-sterilized in 20% bleach for 15 min, rinsed three times with sterile water and germinated on ½-MS media. Rice plants were grown in 10 h light / 14 h dark at 28°C and 26°C, respectively. For heat-stress experiments, 1-week-old rice plants were transferred to a growth chamber at 44°C and 28°C in light and dark, respectively. Rice callus was induced by the method used for rice transformation as previously described^42^.

Tomato plants (*Solanum lycopersicum cv. M82*) were grown under standard greenhouse conditions (16 h supplemental lighting of 88 w/m^2^ at 25°C and 8 h at 15°C). Tomato leaf tissue samples were taken from 2-month-old plants. Tomato fruit pericarp tissues were harvested at 52 days post anthesis (DPA).

### Annotation of LTR retrotransposons

Functional *de novo* annotation of LTR retrotransposons for the genomes of TAIR10 (Arabidopsis), MSU7 (rice) and SL2.50 (tomato) was achieved by the *LTRpred* pipeline (https://github.com/HajkD/LTRpred) using the parameter configuration: minlenltr = 100, maxlenltr = 5000, mindistltr = 4000, maxdisltr = 30000, mintsd = 3, maxtsd = 20, vic = 80, overlaps = “no”, xdrop = 7, motifmis = 1, pbsradius = 60, pbsalilen = c(8,40), pbsoffset = c(0,10), quality.filter = TRUE, n.orf = 0. The plant-specific tRNAs used to screen for primer binding sites (PBS) were retrieved from GtRNAdb^43^ and plantRNA^44^ and combined in a custom fasta file. The hidden Markov model files for gag and pol protein conservation screening were retrieved from Pfam^45^ using the protein domains RdRP_1 (PF00680), RdRP_2 (PF00978), RdRP_3 (PF00998), RdRP_4 (PF02123), RVT_1 (PF00078), RVT_2 (PF07727), Integrase DNA binding domain (PF00552), Integrase zinc binding domain (PF02022), Retrotrans_gag (PF03732), RNase H (PF00075) and Integrase core domain (PF00665). Computationally reproducible scripts for generating annotations can be found at http://github.com/HajkD/ALE.

### ALE-seq library preparation

Genomic DNA was extracted using a DNeasy Plant Mini Kit (Qiagen) following the manufacturer’s instruction. Genomic DNA (100 ng) was used for adapter ligation with 4 µl of 50 µM adapter DNA. After an overnight ligation reaction at 4°C, the adapter-ligated DNA was purified by AMPure XP beads (Beckman Coulter) at a 1:0.5 ratio. *In vitro* transcription reactions were performed using an MEGAscript RNAi kit (Thermofisher) with minor modifications. Briefly, the reaction was carried out for 4 h at 37°C and RNase was omitted from the digestion step prior to RNA purification. Purified RNA (3 µg) was subjected to reverse transcription (RT) using a Transcriptor First Strand cDNA Synthesis Kit (Roche). The custom RT primers were added as indicated for each experiment. After the RT reaction, 1 µl of RNase A/T1 (Thermofisher) was added and the reaction mixture was incubated at 37°C for at least 30 min. Single-stranded first strand cDNA was purified by AMPure XP beads (Beckman Coulter) at a 1:1 ratio and PCR-amplified by 25 cycles using Illumina TruSeq HT dual adapter primers. After purification, the eluted DNA was quantified using a KAPA Library Quantification Kit (KAPA Biosystems) and run on the MiSeq v3 2 X 300 bp platform in the Department of Pathology of the University of Cambridge. The oligonucleotide sequences are provided in Supplementary Table 1.

### Preparation of full-length *Onsen* DNA

The full-length *Onsen* copy (AT1TE12295) was amplified using Phusion High-Fidelity DNA polymerase (New England Biolabs). PCR products were run on 1% agarose gels. The full-length fragment was then purified by QIAquick Gel Extraction (Qiagen) and its concentration measured using the Qubit Fluorometric Quantitation system (Thermo Fisher). Primers used for amplification are listed in Supplementary Table 1.

### RT-qPCR analyses

Samples were ground in liquid nitrogen using mortar and pestle. An RNeasy Plant Mini Kit (Qiagen) was used to extract total RNA following the manufacturer’s instructions. The amount of extracted RNA was estimated using the Qubit Fluorometric Quantitation system (Thermo Fisher). cDNAs were synthesized using a SuperScript VILO cDNA Synthesis Kit (Invitrogen). Real-time quantitative PCR was performed in the LightCycler 480 system (Roche) using primers listed in Supplementary Table 1. LightCycler 480 SYBR green I master premix (Roche) was used to prepare the reaction mixture in a volume of 10 µl. The results were analysed by the ΔΔCt method.

### RNA-seq library construction

Total RNA was prepared as described above. An Illumina TruSeq Stranded mRNA Library Prep kit (Illumina) was used according to the manufacturer’s instructions. The resulting library was run on an Illumina NextSeq 500 machine (Illumina) in the Sainsbury Laboratory at the University of Cambridge.

### Analysis of next-generation sequencing data

For RNA-seq data analysis, the adapter and the low-quality sequences were removed by Trimmomatic software. The cleaned reads were mapped to the MSU7 version of the rice reference genome using TopHat2. The resulting mapping files were processed to the Cufflinks/Cuffquant/Cuffnorm pipeline for quantitation and visualized in an Integrative Genomics Viewer (IGV).

For ALE-seq data analysis, the adapter sequence was removed from the raw reads using Trimmomatic software. For the mapping-based approach, paired-end reads were mapped to the reference genomes (Arabidopsis, TAIR10; rice, MSU7; tomato, SL2.50) using BOWTIE2 with minor optimization (-X 3000). The numbers of reads for each retrotransposon were counted by the featureCounts tool of the SubRead package. IGV was used to visualize the sequencing data. For the alignment-based approach, the forward and reverse reads were merged and converted to fasta files. The fasta files created for all the samples were concatenated and clustered by CD-HIT software with the following options: -c 0.95, -ap 1, -g 1. The resulting fasta file for the representative reads was used as the reference for paired-end mapping. The mapped reads were counted with the featureCounts tool.

For Bisulfite sequencing analysis, raw sequenced reads derived from tomato fruits (52 DPA) and leaves were downloaded from the public repository (SRP008329)^16^ and re-analysed as previously described^46^, with minor modifications. Briefly, high-quality sequenced reads were mapped with Bismark^47^ on the cv. Heinz 1706 reference genome (https://solgenomics.net), including a chloroplast sequence obtained from GenBank database (NC_007898.3) to estimate the conversion rate. After methylation call and correction for unconverted cytosines, the methylation proportions at each cytosine position with a coverage of at least 3 reads were used to generate a bedGraph file for each cytosine context, using the R Bioconductor packages DMRCaller (https://www.bioconductor.org/packages/release/bioc/html/DMRcaller.html) and Rtracklayer^48^. The IGV browser was used to visualize the methylation profiles.

### Detection of retrotransposon insertions

The insertions of selected retrotransposons were detected from the genome resequencing data of *japonica* and *indica* rice accessions downloaded from the 3,000 rice genome project (PRJEB6180). The Transposon Insertion Finder (TIF) program^15^ was used to identify the split reads in the fastq files and detect newly integrated copies. We used MSU7 (http://rice.plantbiology.msu.edu) and ShuHui498 (http://www.mbkbase.org) for *japonica* and *indica* rice, respectively. Only unique non-redundant insertions were considered.

### Data accessibility

The next generation sequencing data generated in this study are deposited in the GEO repository under accession numbers GSEXXXXX.

---

**Supplementary Figure 1.**
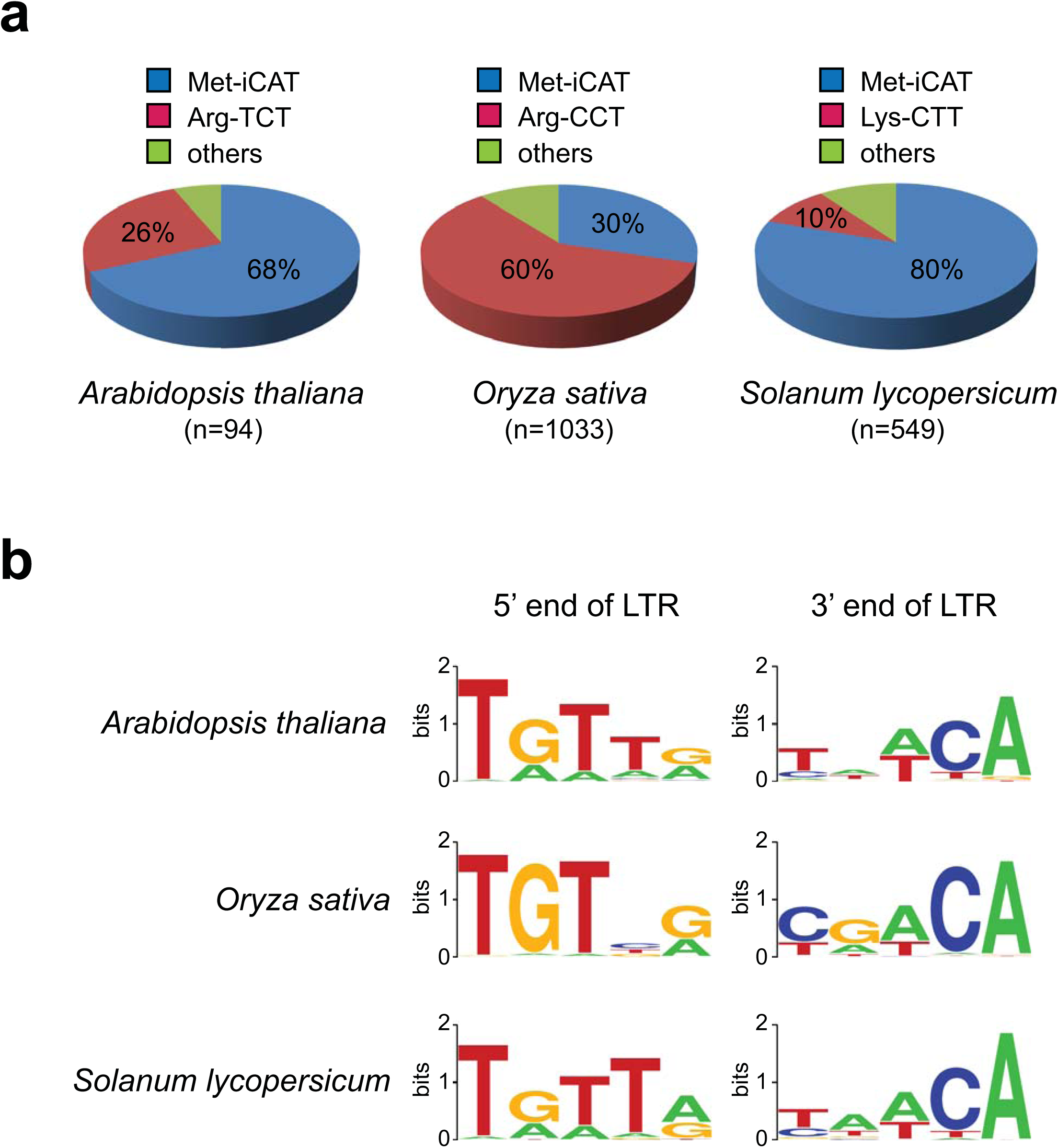
PBS and LTR terminal sequences of LTR retrotransposons in *Arabidopsis*, rice and tomato. **a**, The frequency of tRNAs used for targeting PBS. LTR retrotransposons were annotated by *LTRpred* and selected for young elements by filtering LTR similarities higher than 95%. The total numbers of retrotransposons analysed in each species are shown below. **b**, The conserved sequences of 5’ and 3’ ends of LTR. The first and last five nucleotides of LTRs are displayed. The images were generated by the WebLogo tool (http://weblogo.berkeley.edu/logo.cgi).

**Supplementary Figure 2.**
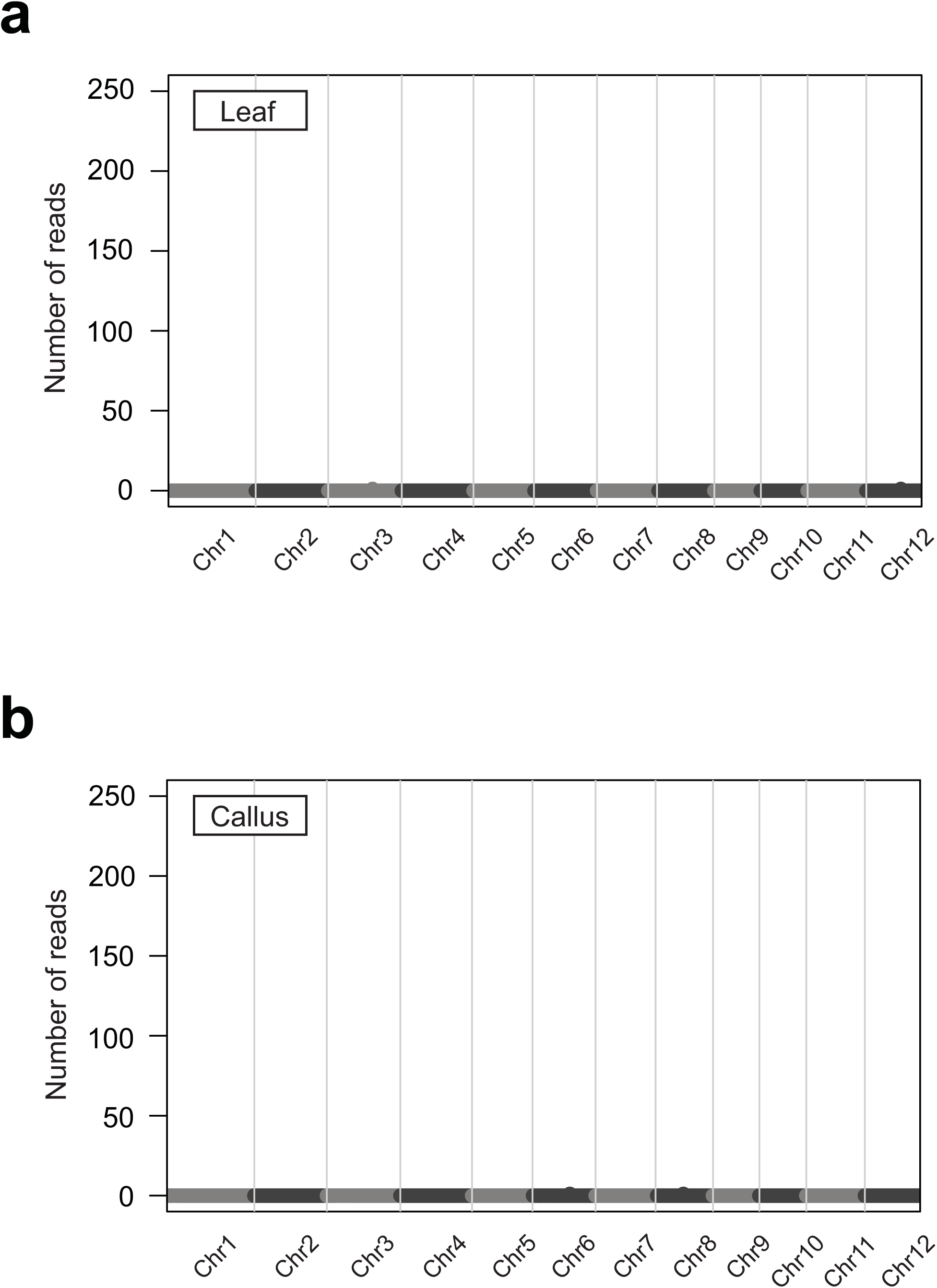
SIRT results from leaves and calli of rice. **a,** Genome-wide plots for SIRT performed in leaves and (**b**) in calli of rice.

**Supplementary Figure 3.**
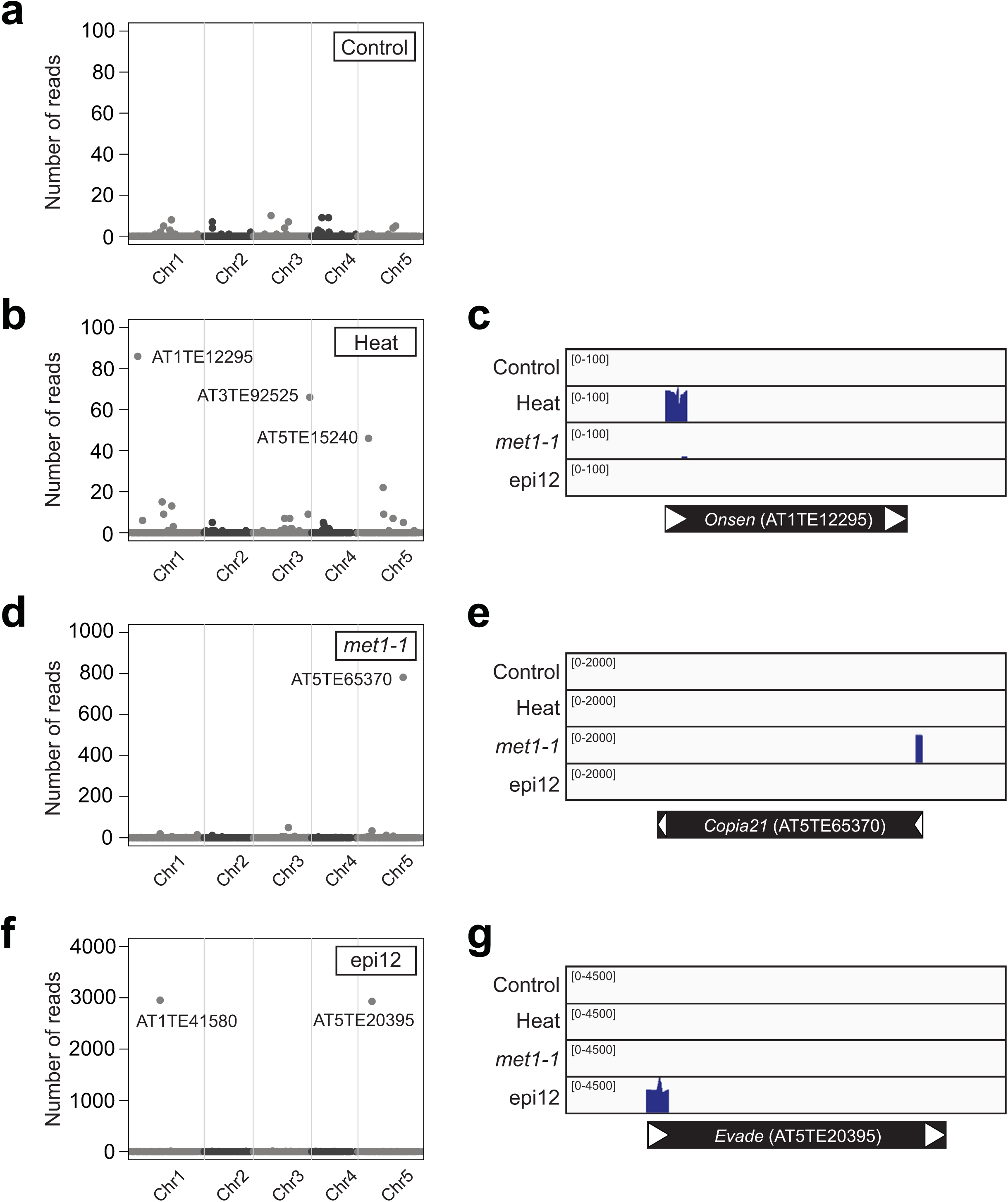
Ale-seq detection of eclDNAs of *Arabidopsis* retrotransposons. Genome-wide plots (**a**, **b**, **d** and **f**) and read coverage plots (**c**, **e** and **g**) for ALE-seq profiles of *Arabidopsis* Col-0 wt (**a**), heat-stressed Col-0 (**b** and **c**), *met1-1* (**d** and **e**), and epi12 (**f** and **g**).

**Supplementary Figure 4.**
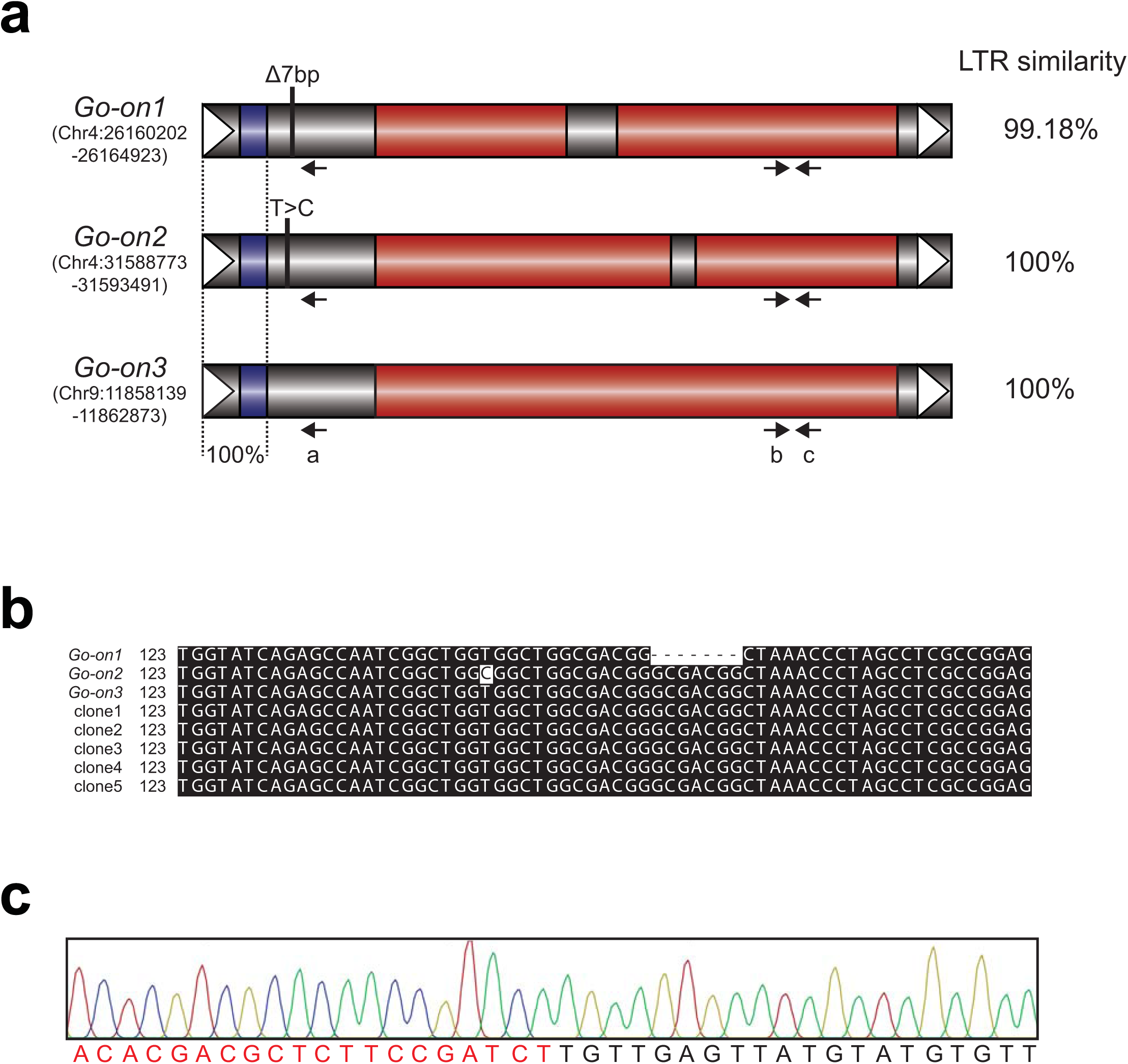
*Go-on* retrotransposon family. **a**, Schematic structure of *Go-on* retrotransposons. The genomic coordinates and LTR similarities of each copy are shown at the left and right, respectively. Red boxes, ORFs; blue boxes, PBS; white arrowheads, LTRs. Note that the sequences of the upstream LTRs through the PBS are identical in all three copies. The sequence variation specific for each element is indicated. Primers used for sequencing and qPCR analyses are shown as arrows. **b**, Multiple sequence alignment of the genomic sequences of three *Go-on* copies and the sequenced ALE clones. ALE-seq was performed using the RT primer specific to *Go-on3* indicated as “a” in **a**. The resulting single-stranded first strand cDNA was PCR-amplified, cloned to the pGEM T-easy vector, and sequenced. Multiple sequence alignment was performed by ClustalW (http://www.genome.jp/tools-bin/clustalw) and visualized by boxshade tools (https://www.ch.embnet.org/software/BOX_form.html). **c**, Sequencing of the ALE-seq product of *Go-on3* showing the junction region of the adapter and LTR. Sequences in red and black are the adapter and *Go-on* LTR, respectively.

**Supplementary Figure 5.**
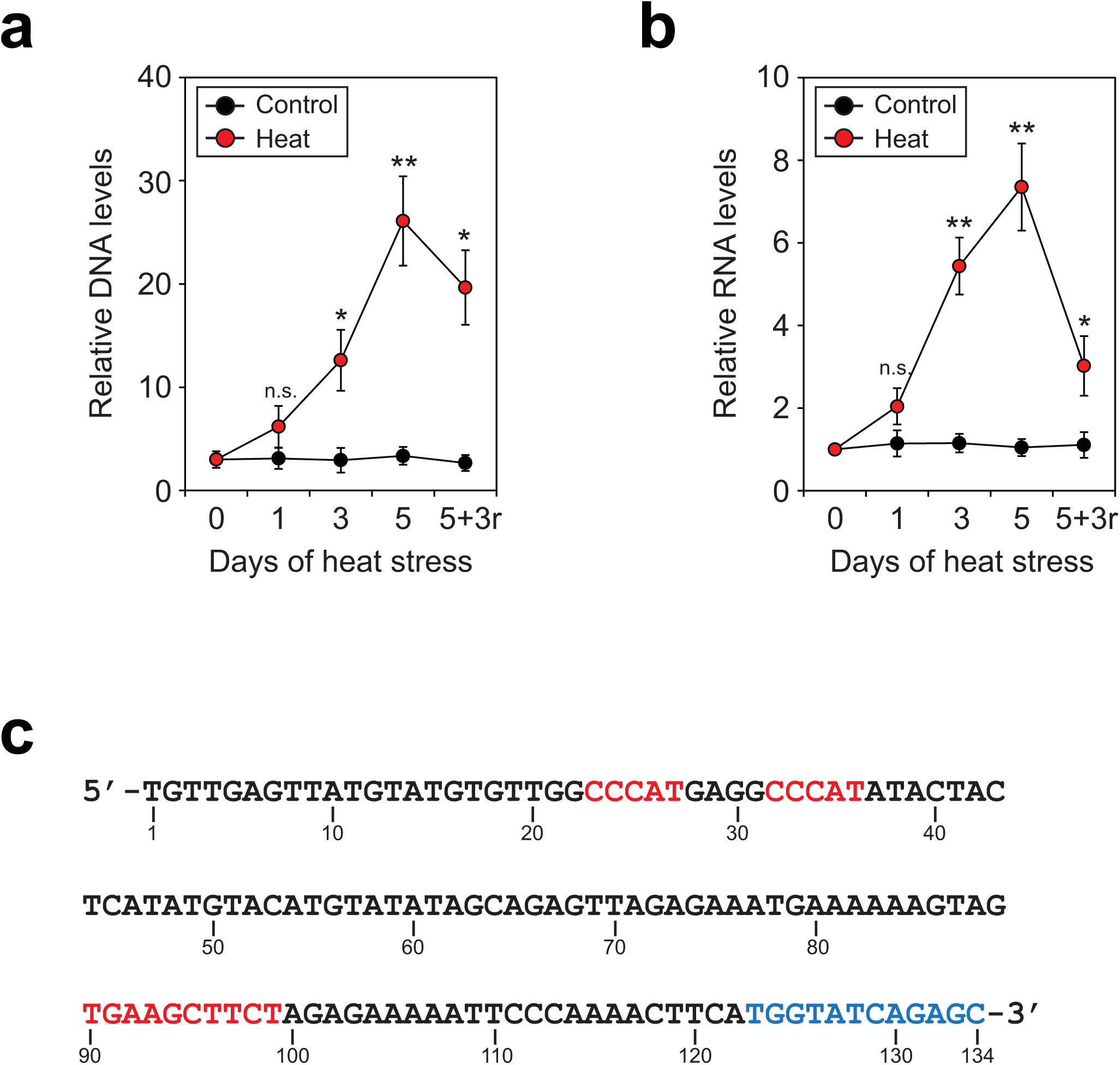
Heat stress-triggered transcriptional activation of *Go-on*. **a** and **b**, The relative levels of DNA (**a**) and RNA (**b**) of *Go-on3* determined by qPCR. Heat treatment (44°C) was applied to 1-week-old rice seedlings for the periods indicated; +3r means 3 days of recovery in normal growth conditions after heat stress. The levels are means ± sd of three biological replicates performed in three technical repeats. For DNA analysis, Day 0 levels are set to 3, reflecting three genomic copies of *Go-on* in *japonica* rice. Normalization was done against *eEF1α*. Asterisks represent significant statistical difference as determined by Student’s t-test. \*\**P* <0.005; \**P* <0.05; n.s., not significant. **c**, The sequence of the left LTR and PBS of *Go-on3*. The sequences in red are the heat-related cis-acting sequence motifs predicted by PlantPan 2.0 tool (http://plantpan2.itps.ncku.edu.tw/index.html). The PBS is shown in blue.

**Supplementary Figure 6.**
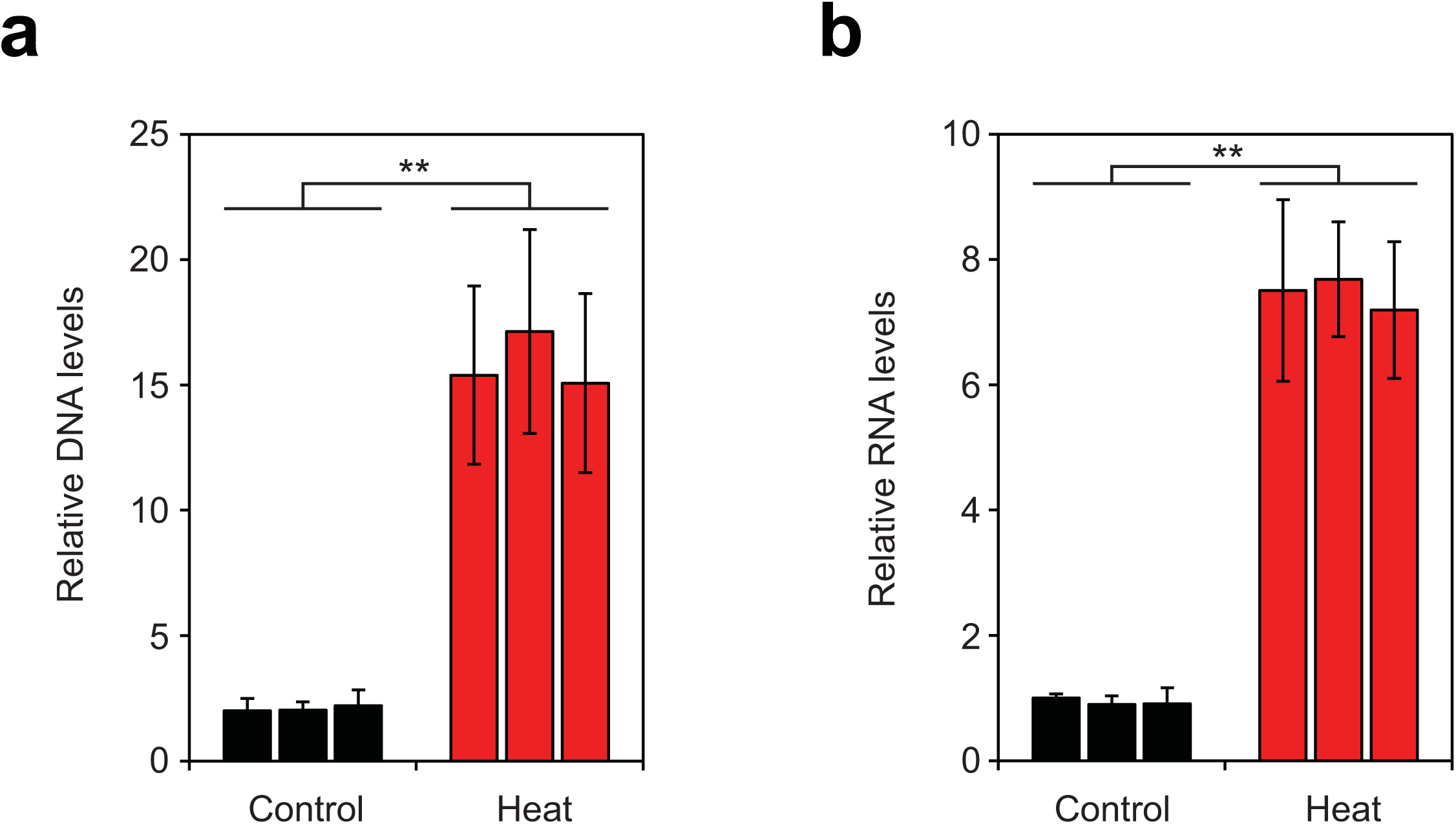
Heat stress-triggered activation of *Go-on* in *indica* rice. **a** and **b**, qPCR analyses for DNA (**a**) and RNA (**b**) levels of *Go-on* in *indica* rice. The levels are means ± sd of three technical repeats. The levels of replicate 1 of control sample are set to 2 (**a**) reflecting 2 genomic copies of *Go-on* in *indica* rice. Asterisks represent significant statistical difference as determined by Student’s t-test. \*\**P* <0.005.

**Supplementary Figure 7.**
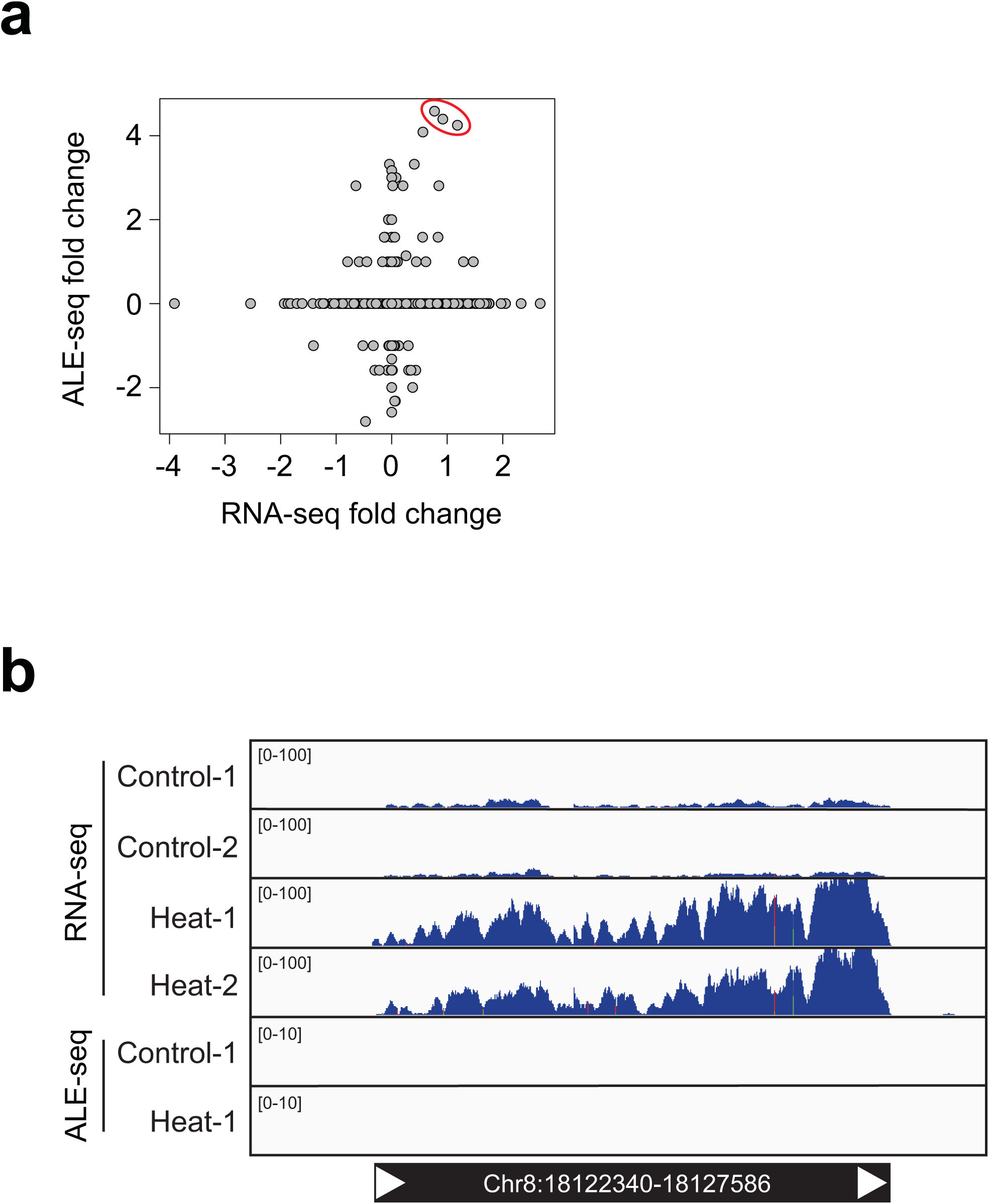
Comparison of mRNA and eclDNA levels. **a**, Scatter plot for fold changes in RNA-seq and ALE-seq profiles in the control and heat-stressed rice plants. Each dot represents an individual retroelement. Three *Go-on* retroelements are circled in red. **b**, Read coverage plot for a selected retrotransposon showing evidence of transcriptional activation upon heat stress not followed by synthesis of eclDNAs.

